# Relevance of the viral Spike protein/cellular Estrogen Receptor-α interaction for endothelial-based coagulopathy induced by SARS-CoV-2

**DOI:** 10.1101/2022.10.04.510657

**Authors:** Silvia Barbieri, Franca Cattani, Leonardo Sandrini, Magda Maria Grillo, Carmine Talarico, Daniela Iaconis, Lucia Lione, Erika Salvatori, Patrizia Amadio, Gloria Garoffolo, Mariano Maffei, Francesca Galli, Andrea Rosario Beccari, Emanuele Marra, Marica Zoppi, Michael Michaelides, Giuseppe Roscilli, Luigi Aurisicchio, Riccardo Bertini, Marcello Allegretti, Maurizio Pesce

**Author notes:** First equal contribution (Silvia Barbieri, Franca Cattani). Last equal contribution (Marcello Allegretti, Maurizio Pesce). Correspondence to: Maurizio Pesce, PhD - Unità di ricerca in ingegneria tissutale cardiovascolare; Silvia Stella Barbieri, PhD - Unità di ricerca asse cuore cervello: meccanismi molecolari e cellulari. Centro cardiologico Monzino, IRRCS. Via C. Parea, 4, I-20138, Milano, Italy.

## Abstract

Severe coagulopathy has been observed at the level of the microcirculation in several organs including lungs, heart and kidneys in patients with COVID-19, and in a minority of subjects receiving the SARS-CoV-2 vaccine. Various mechanisms have been implicated in these effects, including increases in circulating neutrophil extracellular traps, excessive inflammation, and endothelial dysfunction. Even if a correlation between infection by SARS-CoV-2 and upregulation of coagulation cascade components has been established in the lung, no direct proofs have been yet provided about the transcriptional machinery controlling the expression of these factors. Recent results obtained by us reported a novel transcriptional function of the SARS-CoV-2 Spike (S) viral protein involving a direct protein-protein interaction with the human Estrogen Receptor-α (ERα). Given the implications of ERα in the control of key effectors in the coagulation cascade, we hypothesized that S-protein might increase the pro-coagulation activity of endothelial cells *via* the transcriptional activity of the ERα, thus justifying the enhanced risk of thrombosis. To assess this, we tested the effects of S-protein on the expression of Tissue Factor (TF) and the overall procoagulation activity in a human endothelial cell line and confirmed this finding by overexpressing S-protein by gene transfer in mice. We then designed and tested two-point mutations in the S2 S-protein sequence that abolished the pro-coagulation function of S-protein *in vitro* and *in vivo*, without compromising its immunogenicity. In addition to reveal a new potential transcriptional function of S-protein, these results inspire the design of new vaccines with lower risk of thrombogenesis. Indeed, while the benefit/risk ratio remains overwhelming in favor of COVID-19 vaccination, our results shed light on the causal mechanisms of some rare anti-SARS-CoV-2 vaccine adverse events, and are thus essential for current and future vaccination and booster campaigns.

## Introduction

The propensity to develop diffuse and severe coagulation in the microcirculation in numerous organs, leading to multi-systemic organ failure, is one of the life-threatening conditions characterizing the acute infection by SARS-CoV-2 (*1*, *2*). Several mechanisms may account for the hyper-coagulatory conditions observed in COVID-19 patients. Indeed, SARS-CoV-2 induces endothelial inflammation/dysfunction (*3*), platelet hyper-reactivity (*4*), monocyte activation (*5*), and generation of neutrophil extracellular traps (*6*), all of which promote and support the activation of the coagulation cascade in an infection-dependent manner. Accordingly, increased levels of tissue factor (TF), the main initiator of coagulation, have been found in tissues and in the circulation of COVID-19 patients (*7*).

The outcome of the COVID-19 vaccination campaign has also revealed a potential risk of vaccine-induced immune thrombotic thrombocytopenia (VITT) or thrombosis with thrombocytopenia syndrome (TTS) (*8*, *9*). Although this risk is extremely low - in the range of few hundred or lower number of cases per million of vaccinated subjects (*10*), it has called for caution especially considering the worldwide impact of the SARS-CoV-2 vaccination campaign with DNA or RNA vaccines, and the necessity to evolve stratification criteria to identify subjects with a potential risk for thrombosis in response to DNA- or RNA-based vaccination against this and, potentially, other coronaviruses (*11*). By using bioinformatics and the EXaSCale smArt pLatform Against paThogEns (EXSCALATE) supercomputing platform (Solis et al., *Sci. Adv*. in press) of potential cellular interactors of the S-protein, we predicted a potential function of the viral protein as a cofactor for ERα nuclear signaling. This function arises from the direct interaction of a nuclear receptor coregulator (NRC) LXD-like motif present on the S2 subunit of the viral protein, and the activation function 2 (AF-2) region on ERα. In the present work we provide evidence that the S-protein increases the procoagulation activity, and upregulates TF expression/activity in human endothelial cells *via* the ERα signaling pathway. To validate these findings, we used selective estrogen receptor modulators (SERMs) and selective estrogen receptor degraders (SERDs) Raloxifene and Fulvestrant/ICI 164,384 (*12*), and designed an expression vector that drives a robust production of S-protein in mice based on DNA electroporation into the adductor muscle. Finally, based on our previous molecular interaction modelling (Solis et al., *Sci. Adv*. in press), we designed two mutated versions of the S-protein that were predicted to lose the interaction with ERα, and that experimentally lacked pro-coagulatory activity *in vitro* and *in vivo*.

## Materials and methods

### 3D Proteins Selection and molecular dynamics simulation workflow

The S-protein 3D model was built based on the experimental coordinates deposited on the Protein Data Bank database with code 6VYB returned to its wild-type form and fully glycosylated (*13*). An asymmetric glycosylation of the three protomers has been derived by glycoanalyitic data for the N-glycans and O-glycans as previously reported (*14*). For the estrogen receptor, the X-RAY PDB model with code 3UUD was used, containing ER_α_ and Nuclear receptor coactivator-2 (*15*) and 3OLL, containing ER_β_ and Nuclear receptor coactivator-1(*14*). The proteins were modeled using Amber14SB force field (*16*), the carbohydrate moieties by the GLYCAM06j-1 version of GLYCAM06 force field (*17*), and the general amber force field (GAFF) was used for the estradiol bound to ER receptor. The so prepared structure was used as starting point for MD simulations. Protein was inserted in a clinic box, extending up to 10 Å from the solute, and immersed in TIP3P water molecules. Counter ions were added to neutralize the overall charge with the genion GROMACS tool. After energy minimizations, the system was relaxed for 5 ns by applying positional restraints of 1000 kJ mol^-1^ nm^-2^ to the protein atoms. Following this step, unrestrained MD simulation was carried out with a time step of 2 fs, using GROMACS 2020.2 simulation package (supercomputer Marconi-100, CINECA, Bologna, Italy) V-rescale temperature coupling was employed to keep the temperature constant at 300 K. The Particle-Mesh Ewald method was used for the treatment of the long-range electrostatic interactions. The first 5ns of each trajectory were excluded from the analysis. The trajectory obtained after 1 microsecond MD simulation has been clustered in order to obtain representative structures. In particular, the structure used for the docking studies is the first centroid of the first cluster extracted from the MD experiment. For the ER, the XRAY PDB model with code 3OLL was used, containing 17_β_-Estradiol and Nuclear receptor coactivator 1 (*18*).

### S-ERα Protein-Protein Docking procedure

The input of two individual proteins were set up. In particular, the S-protein and ER were used as receptor and ligand respectively. Then, the HDOCK tool was ran to sample putative binding modes through an FFT-based search method and then scoring the protein–protein interactions. Then, the top 100 predicted complex structures are produced as output, and the best ten hypotheses were visually inspected to confirm the reliability of the calculation. The entire workflow is well described in the work published by Yan et al. (*19*).

### Manufacturing of S-protein, Sp5 and Sp7 for in vitro testing

Manufacturing of S-protein and its mutants (Sp5 and Sp7) in HEK293 cell expression system was based on the wild-type S-protein sequence (*20*) deleted from the transmembrane domain (aa 17-1213) and containing stabilizing mutations. For Sp5 mutant protein generation the sequence LGDIA was muted in AGDAA, whereas for Sp7 version the sequence LPPLL was muted in APPAA. The nucleotide sequences encoding the trimeric S-protein and the different mutants were then generated and inserted into a plasmid DNA vector. This vector encompassed the Human cytomegalovirus (CMV) immediate early enhancer and promoter, the murine Ig kappa chain leader sequence for protein secretion, the bovine growth hormone polyadenylation (bGH-PolyA) signal, a specialized termination sequence for protein expression in eukaryotic cells and the kanamycin resistance gene from *Staphylococcus aureus* for plasmid amplification in bacteria.

The trimeric S-protein and its mutants were produced by transient transfection of Expi293F high-density cells with the ExpiFectamine 293 (Thermo Fisher) lipid cationic transfection reagent according to the manufacturer’s instructions. The supernatant containing the proteins was collected after five days of incubation from start of transfection and subjected to clarification by centrifugation and filtration for the subsequent purification steps. The proteins were batch purified using IMAC (Immobilized Metal Chelate Affinity Chromatography) using PureCube Ni-NTA Agarose resin slurry (Cube-Biotech). Briefly, the resin slurry was centrifuged and incubated with equilibration buffer (50mM NaH2PO4, 500mM NaCl, pH 7.4, 10mM imidazole). The equilibrated resin was combined with the culture supernatant containing the recombinant proteins and incubated o/n, 4°C, on a rotating platform. The resin was subsequently collected by centrifugation, washed, and the protein was eluted by an elution buffer containing 300 mM Imidazole and subjected to dialysis in phosphate buffer (PBS) using slide-A-lyzers (Thermo Fisher) as indicated in the product datasheet. Once recovered from dialysis, the proteins were quantified on a spectrophotometer measuring the absorbance at 280 nm. Trehalose (Sigma-Aldrich) was used (5% w/v) as a stabilizer of *in vitro* produced S-proteins (*21*). The purity of the proteins was assessed by SDS-PAGE and Western Blot analysis, conducted under both reducing and non-reducing conditions. Furthermore, the functional activity linked to correctness of the trimeric organization was demonstrated by the binding to the human ACE-2 receptor (Octet Red-Forte Bio).

### Cells and treatment

EA.hy926 cells (ATCC) were cultured in complete medium: DMEM supplemented with penicillinstreptomycin, non-essential amino acids (NEAA, ThermoFisher Scientifics), tricine buffer (Sigma-Aldrich), HAT (Sigma-Aldrich), and 10% fetal bovine serum (FBS, HyClone, Thermo Scientific) on 0.2% gelatin-coated plates. Cells were seeded at a 150.000 cells/ml density in complete medium, and at confluence (after about 48 hours) were starved for 24 hours in fresh DMEM supplemented with 2% FBS. After starvation they were cultured for 48 hours in DMEM 0.5% FBS with TNFα (50 ng/ml, Peprotech), 17β-Estradiol (10 - 100 nM; Sigma-Aldrich), Raloxifene (2 μM; Sigma-Aldrich), Fulvestrant (100 nM; Sigma-Aldrich) and/or wildtype S-protein and, and the two Sp5/Sp7 mutants. Trehalose (the protein stabilizer employed in S-protein, Sp5 and Sp7 preparation) was tested in preliminary experiments to assess possible toxic effects. In no case the addition of this compound to culture medium determined cell death (data not shown).

### Quantitative real-time polymerase chain reaction (qRT-PCR)

For qRT-PCR experiments, cells were harvested in TRIzol Reagent (ThermoFisher Scientifics) according to the manufacturer’s instructions. 1 μg of RNA was reverse transcribed using iScript^™^ Advanced cDNA Synthesis Kit (Biorad). qRT-PCR was then carried out to assay TF (TF: forward 5’-CCCAAACCCGTCAATCAAGTC-3’, reverse 5’-CCAAGTACGTCTGCTTCACAT-3’) and 18S ribosomal RNA (18S: forward 5’-CGGCTACCACATCCAAGGAA-3’, reverse 5’-CCTGTATTGTTATTTTTCGTCACTACCT-3’) used as internal reference genes. Samples of cDNA (2.5 μL) were incubated with 25 μL of containing TF or 18S primers and fluorescent Luna^®^ Universal qPCR Master Mix (New England Biolabs), and qRT-PCR was carried out in triplicate for each sample on the CFX Connect real-time System (Bio-Rad Laboratories).

### Procoagulant activity

Samples were lysed with 15 mM n-Octyl-B-D-glucopyranoside lysis buffer at 37°C for 10 min, sonicated at 20 kHz for 20 seconds and diluted with 25 mM HEPES saline. The total protein concentrations were determined using the Bradford method. 60 μl of homogenate (0.07 mg/mL) were mixed with 60 μL citrated pooled human plasma and 60 μL CaCl_2_ (final concentration 25 mM), and procoagulant activity was quantified by a one-stage plasma recalcification time assay. Clotting times were expressed in relative U/μg protein based on a standard curve of serially diluted human thromboplastin preparation.

### TF activity

ACTICHROME® TF activity assay was measured according to manual instructions (BioMedica Diagnostics). Briefly, Briefly, cells were lysed in a recommended buffer (50mM Tris-HCl, 100mM NaCl, 0.1% Triton X-100, pH 7.4), sonicated and incubated at 37°C for 30 min. TF activity was detected by two stage chromogenic assay interpolating the mean absorbance values of test samples, directly from the standard curve.

### In vivo gene transfer and relative S-protein dosage/immunization

The study was carried out in accordance with UKCCCR guidelines for the welfare of animals as well as the European Directive 2010/63/EU. For *in vivo* expression and immunogenicity assessment, 6-8 weeks old C57BL/6 mice (n=10-16/group, Envigo, USA) were injected intramuscularly (i.m.), particularly in the quadriceps, with either DNA plasmid or amplicon (dose ranging from 10 μg to 50 μg) and electrically stimulated as previously described (*22*). The DNA was formulated in Phosphate Buffered Saline (PBS). DNA-EP was performed by means of a Cliniporator Device EPS01 and using N-10-4B electrodes (IGEA, Italy) with the following electrical conditions in Electro-Gene-Transfer (EGT) modality: 8 pulses 20 msec each at 110V, 8Hz, 120msec interval. Mice with electroporation procedure only were used as control group. After 24, 48 and 96 hours after electroporation, animals were anesthesized and blood collected by cardiac venipuncture into 3.8% sodium citrate. For plasma preparation citrated blood was centrifuged (within 30 minutes) at 3000 rpm for 20 min, and immediately stored in dry ice and then at −80 °C until analysis of pro-coagulatory activity.

### Plasma Clotting Time

Recalcified plasma clotting times was evaluated according to a method previously described (*23*). Briefly, to 30 μL mouse platelet-poor plasma (PPP) were added 30 μL of HEPES saline buffer and 20 μL of citrated PPP obtained from pooled wild-type mouse. Samples, pre-wormed for 1 minute in the thermostatically-controlled water bath at 37°C, were mixed to 40 μL of CaCl_2_ 25mM and incubated at 37°C with shaking until clot formation. The plasma clotting time was measured as the time it takes for the plasma to undergo gelation, detected by loss of movement of the plasma in response to the rotation and shaking by an operator blinded to the experimental groups.

### Immunogenicity determination by ELIspot and serum titration

For the ELIspot a commercial kit (Mabtech) was used. Briefly, splenocytes collected from mice inoculated with the DNA vectors 35 days earlier were plated at 250,000 and 500,000 in duplicate for each condition to be tested (DMSO, Pool S1, Pool S2, and Concanavaline A). The S1 and S2 peptides used as an overnight stimulus were used at a concentration of 1μg/ml; concanavaline A (C5275, Sigma) was used as a positive control at a concentration of 250 μg/ml. Results were reported as SCF (Spot Forming Cells)/10^6^ splenocytes. For sera titration an in-house ELISA assay was developed. Briefly, full length wild-type S protein was coated on a 96well plate at a concentration of 1μg/ml in PBS in a final volume/well of 50 μl. The plate was incubated overnight at 4°C and after 5 washes with PBS1X-Tween 0,05% was blocked with 150ul of BSA3% in PBS1X-Tween 0,05% 1h at RT in agitation. Scalar dilutions of sera collected from transfected mice at day 35 (1:300; 1:900 1:2700 1:8100 1:24300 1:72900 1:218700) were then incubated overnight at 4°C. After 5 washes with PBS1X-Tween 0,05% an anti-mouse IgG (H + L)-HRP Conjugate (Biorad) diluted 1:2000 in PBS1X-Tween 0,05% was added for 1h at RT. Plate was developed with 50ul of Alkaline Phosphatase Yellow (pNPP, Sigma) for 30 minutes, and read at 405 nM in a Microplate reader (Tecan). Endpoint titer was calculated by plotting the log10 OD and the log10 sample dilution. A regression analysis of the linear part of the curve allowed calculation of the endpoint titer. An OD of 0.2 was used as a threshold.

### Statistical analyses

Statistical treatment of the data was performed using the Graph (Prism) software (version 9.0). Bar graphs were generated using the same program. The choice of statistical tests was done based on the possibility to perform paired/unpaired statistical tests using parametric/non-parametric tests following data normality checking with Kolmogorov-Smirnov test. The type of tests used as well as the level of significance is indicated in the figure legends and the figures, respectively. A *P* < 0.05 was considered an acceptable significance level to assess a statistically-validated difference among treatments/samples.

## Results and Discussion

In order to clarify the relevance of S-protein for the expression/release of pro-coagulation factors by the endothelium, we performed *in vitro* experiments using the immortalized human EC line Ea.Hy926. We exposed these cells to increasing concentrations of the S-protein and, as controls, 17β-Estradiol (ES), a natural ERα agonist (*24*), TNFα a pro-inflammatory cytokine involved in SARS-CoV-2 coagulopathy (*25*), Raloxifene (RAL), a SERM (*26*), and Fulvestrant (FS), a SERD (*27*). The results of these experiments are reported in **Figure 1**, where we show that the exposure to incremental amounts of the S-protein increased the pro-coagulant activity (PCA) of the cells (**Figure 1a, c**) and the expression of *TF* at a mRNA level (**Figure 1b, d**). It is important to note that the S concentrations used here were in line with the reported level of circulating Spike in COVID-19 patients (*28*), and well above the maximal values attained after inoculating both a first and a second dose of the currently available mRNA vaccines - a concentration ≤0.1 ng/ml according to Ogata et al. (*29*). This difference might explain the larger risk of coagulative adverse events in the course of COVID-19 infection than after vaccination. The use of both inhibitors of ERα signaling significantly (although not totally) blunted the effect of S-protein, suggesting an involvement of the S-protein/ERα interaction in the control of the pro-coagulation activity in endothelial cells. It is important to note that an almost complete inhibition of PCA and mRNA expression of *TF* by both ERα inhibitors was observed in a range of S-protein concentrations (1-100 ng/ml) superimposable to the one reported in the serum of COVID-19 patients for non-infective pathological effects of SARS-CoV-2 (*28*).

**Figure 1.**
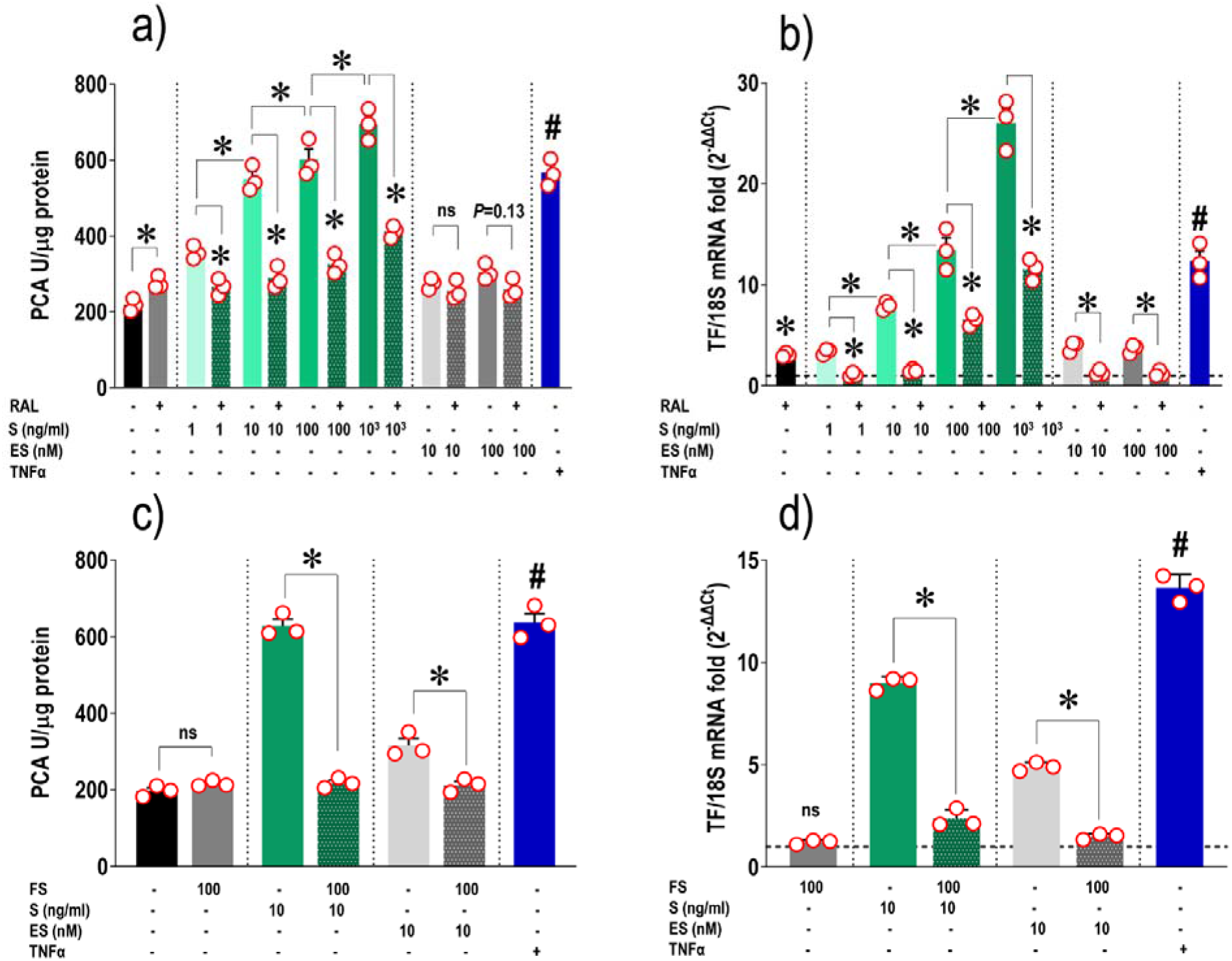
Pro-coagulatory activity (PCA) and effect on *TF* mRNA expression of increasing doses of S-protein in ECs; inhibition effects of S-protein activity by Raloxifene (RAL) and Fulvestrant (FS). Panels **a** and **b** indicate, respectively, the PCA and the fold changes in *TF* mRNA expression (expressed as 2^-ΔΔCt^) in ECs stimulated with increasing doses of wild-type S-protein (1 - 1000 ng/ml) ± Raloxifene, 17β-Estradiol (ES) ± Raloxifene, and TNFα. Panels **c** and **d** show the PCA trend and the *TF* mRNA expression after treating cells with S-protein (10 ng/ml) and ES ± Fulvestrant. Note that both Raloxifene and Fulvestrant reduced significantly the pro-coagulatory activity and the expression of *TF* in ECs, strongly suggesting the relevance of the S-protein/ERα interaction for activation of the coagulation cascade. * indicate P < 0.05 as calculated by repeated measures one-way ANOVA with Tukey multiple comparisons *post-hoc* tests. # indicate the same level of significance in the comparison between control cells and cells treated with TNFα, used as a positive control. Statistical comparisons for data contained in panels **b** and **d** were performed using the ΔCt values to include controls (equalized to 1 and indicated by the dotted line in both fold change graphs). The number of experimental replicates included in the analysis is *n* = 3 and it is indicated by the individual circles overlapped to graphs.

In a recent research work aimed at characterizing the interaction of S-protein with ERα, we identified in the S2 domain putative LXD-like motifs potentially mimicking Nuclear Receptor Coactivator (NCOA) function (Solis et al., *Sci. Adv*. in press). *In silico* molecular protein-protein docking simulations supported a S-protein/ERα binding mode hypothesis. Molecular Dynamics (MD) simulations of the best docking complexes were carried out, and this showed the formation of a strong interaction between ER and S-protein even in the initial stages of the protein-protein recognition. The best obtained binding hypothesis highlighted the binding of ER to the S-protein region containing the two described LXD-like motifs, red and light blue surfaces in **Figure 2a**.

**Figure 2.**
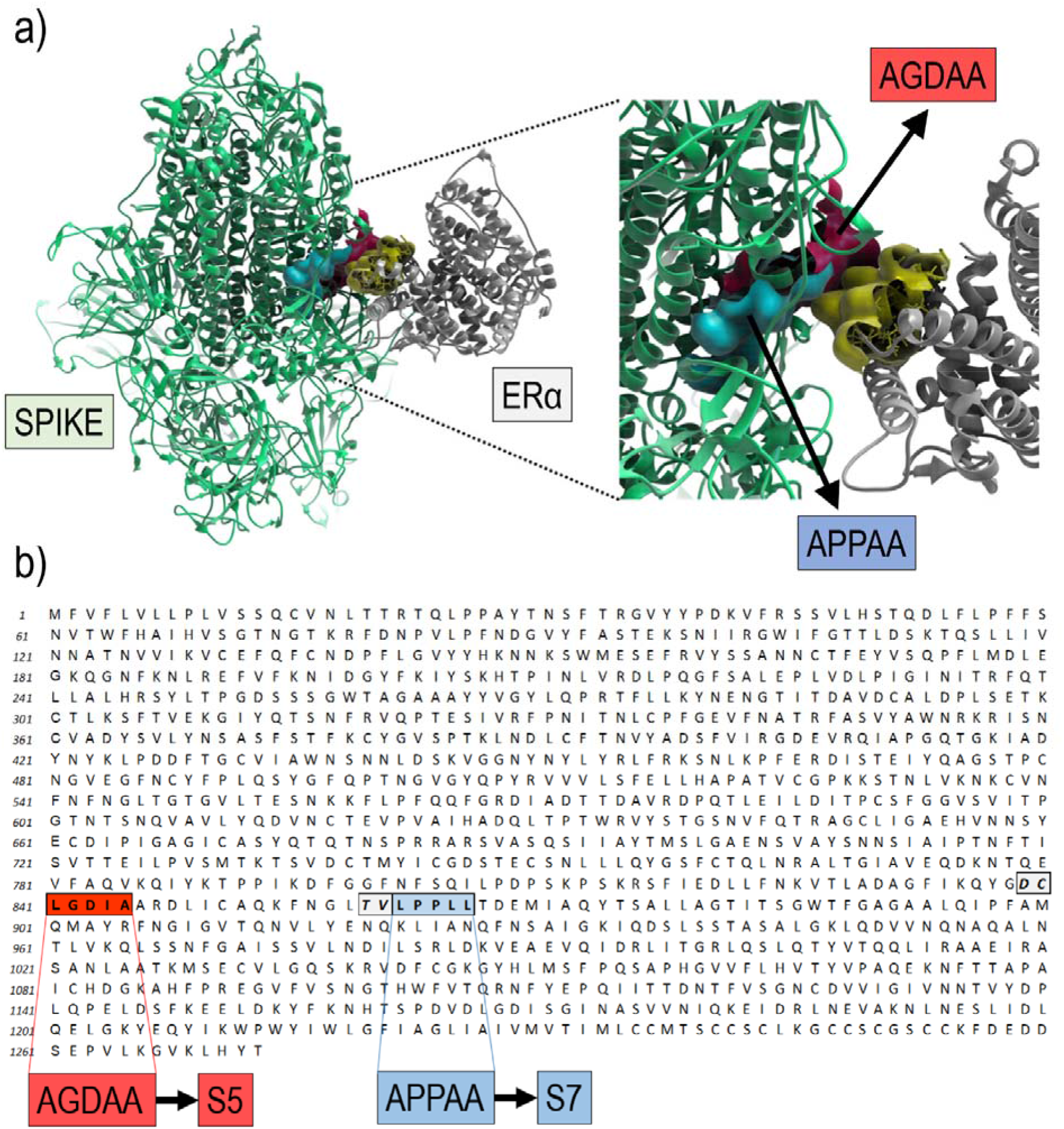
Computer modeling of the S-protein/ERα interactions at conserved LXD nuclear receptor coregulatory (NRC) motifs, and S-protein mutants design. **Panel a**, shows the computer-modeled S-protein/ERα proteins structures and interaction. The region (helix-12) of the ERα interacting with S-protein is highlighted in yellow, while the two LDX motifs in S-protein are indicated as red and light blue surfaces, respectively. Sp5 mutation lies in the red region, whereas Sp7 is in the light blue surface. **Panel b** shows the primary aminoacidic sequence of S-protein with an indication of two LXD-like patterns evidenced in the same color as in **panel a**. The LXXLL motif and a homologous region are highlighted in red (S5) and blue (S7) boxes, respectively. −1 and −2 positions are reported in italic and light grey background.

The knowledge gained based on MD simulation guided us in introducing point mutations in these regions that might potentially abolish the interaction, and thus minimize the intracellular biological functions of the S-protein/ERα complex. To this aim, starting from structural domains involved in S-protein/ERα interactions identified by using bioinformatics and the EXSCALATE platform EXaSCale smArt pLatform Against paThogEns (EXSCALATE) (*30*), we selected two key regions with the (LxxLL) protein sequence (**Figure 2a, b**) and designed the two S-protein mutated versions, named Sp5 and Sp7 (see materials and methods, **Figure 2b**) predicted to lack affinity to ERα while maintaining the wild-type conformational structure. It is important to note that the two LXD-like patterns reported in the primary aminoacidic sequence of S-protein **(Figure 2b**), are not related to known amino acid mutations reported in SARS-CoV-2 variants (see: https://gisaid.org/; https://covariants.org/shared-mutations). S-protein, Sp5 and Sp7 were manufactured as recombinant proteins in HEK293 cell expression system and after purification, to confirm the wild-type folding, they were characterized by affinity assay toward the hACE2 receptor. The obtained K_D_ (M) values (S-protein: 6.4×10^-10^ ± 5.9×10^-12^; Sp5: 2.6×10^-10^ ± 4.7×10^-12^; Sp7: 4.1×10^-10^ ± 5.1×10^-12^) confirmed the equivalence of the three proteins in maintaining the wild-type conformational structure. In order to validate the biological activity of these two mutated proteins, we first assessed the effect of the Sp5 and Sp7 trimeric S-protein mutants on the ECs PCA and TF expression. The results showed a marked difference in the response of cells to exposure to the two mutants compared with the wild-type S-protein with a significant reduction of the PCA and only a slight upregulation of *TF* transcript (**Figure 3 a, b**). This finding was confirmed by analysis of the cellular TF activity, which showed that treatment with the two mutants prevented TF activation (**Figure 3c**). It was interesting to note that while the pro-coagulatory effect of the two mutants was not reduced at the level of the control cells (comparisons of light brown and light purple bars *vs*. grey bar in **Figure 3a** rendered both significant differences according to the employed statistics), suggesting that other S-protein related mechanisms account for a residual activity, the activity of TF was entirely blunted by the mutants, in line with findings obtained with the antagonists with SERMs or SERDs characteristics.

**Figure 3.**
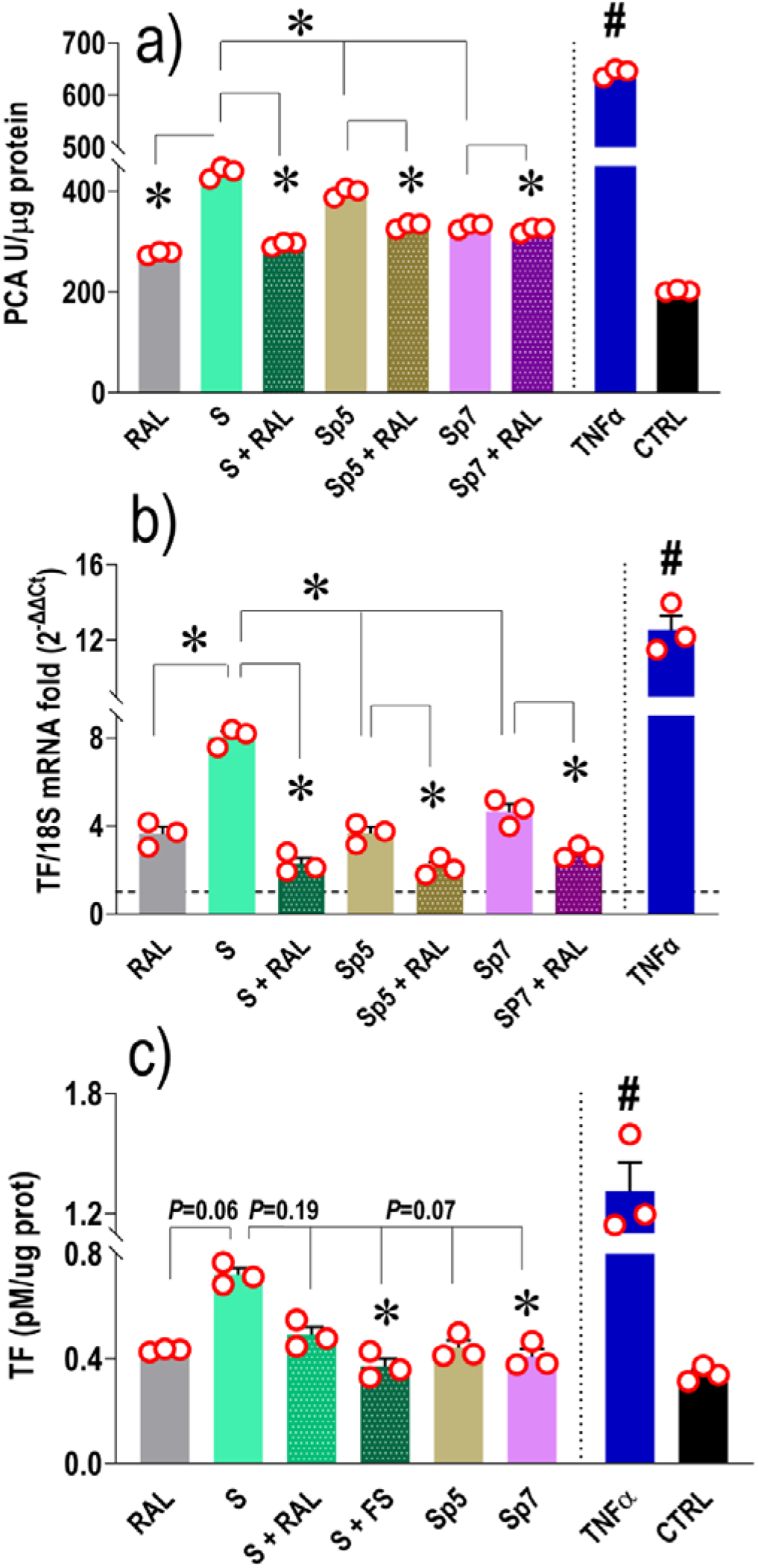
Pro-coagulatory activity and TF mRNA/activity levels in ECs treated with wild-type Sprotein,Sp5 and Sp7 mutants. Panels **a** and **b** indicate that Sp5, and to a higher extent Sp7 mutants, do not elicit a strong coagulative response in ECs. This finding is confirmed by a lower trend in presence of active TF in ECs extracts comparable to that observed in untreated cells and cells treated with wild-type S-protein in the presence of both Raloxifene and Fulvestrant (panel **c**). Statistical comparisons for data contained in panels **b** were performed using the ΔCt values to include controls (equalized to 1 and indicated by the dotted line in the fold change graphs). * indicate *P* < 0.05 as calculated by repeated measures one-way ANOVA with Tukey multiple comparisons post-*hoc* tests. # indicate the same level of significance in the comparison between control cells and cells treated with TNFα, used as a positive control. The number of experimental replicates included in the analysis is *n* = 3 and it is indicated by the individual circles overlapped to graphs.

The S-protein coding sequences (wild-type, and the two Sp5/Sp7 mutants, **Figure 2**) were introduced into a proprietary vector (pTK1A) under the control of human CMV/intA promoter to elicit a strong transgene production in line with the existing expression systems employed in SARS-CoV-2 vaccination (*22*, *31*). To increase the level of transcription, an optimal translation initiation (Kozak) sequence was inserted upstream of the ATG. Two consecutive stop codons were inserted downstream of the codon sequence, followed the bovine growth hormone (bGH) polyadenylation signal. The cDNA molecular design was based on a bioinformatic study and according to existing studies (*32*). Codon-optimized S-protein mutants were synthesized taking into account codon usage bias, GC content, CpG dinucleotides content, mRNA secondary structure, cryptic splice sites, premature PolyA sites, internal chi sites and ribosomal binding sites, negative CpG island, RNA instability motif (ARE), repeat sequences (direct repeat, reverse repeat and Dyad repeat) and restriction sites that may interfere with cloning. In addition, to improve translational initiation and performance Kozak and Shine-Dalgarno Sequences were inserted in the synthetic genes. To increase the efficiency of translational termination, two consecutive stop codons were inserted at the end of cDNAs. The codon usage bias in human was upgraded to a CAI of at least 0.94. GC content and unfavorable peaks were optimized to prolong the half-life of the mRNA. The wild-type S, Sp5 and Sp7 DNA vectors were amplified, and endotoxin-free preparations were inoculated in the lower limb adductor muscles in mice to assess pro-coagulatory activity and at early times (48, 96hrs) after administration. As shown in **Figure 4a**, inoculation of the vector carrying the wild-type S-protein significantly shortened the clotting time at 48hrs, but not at 96hrs. This indicates that the *in vivo* expression of the S-protein causes a transient increase in the blood clotting followed by a relapse. This confirms the *in vitro* data and might explain the rapid increase in coagulopathy observed in patients with COVID-19 (*7*) and, in rare cases, the hypercoagulopathy observed after administration of the vaccines (*2*, *33*). Interestingly, the administration of the vectors carrying the two mutant proteins did not have the same effect, suggesting that the two mutations preventing the interaction with ERα do not affect the coagulation cascade like the wild-type S-protein (**Figure 4a**).

**Figure 4.**
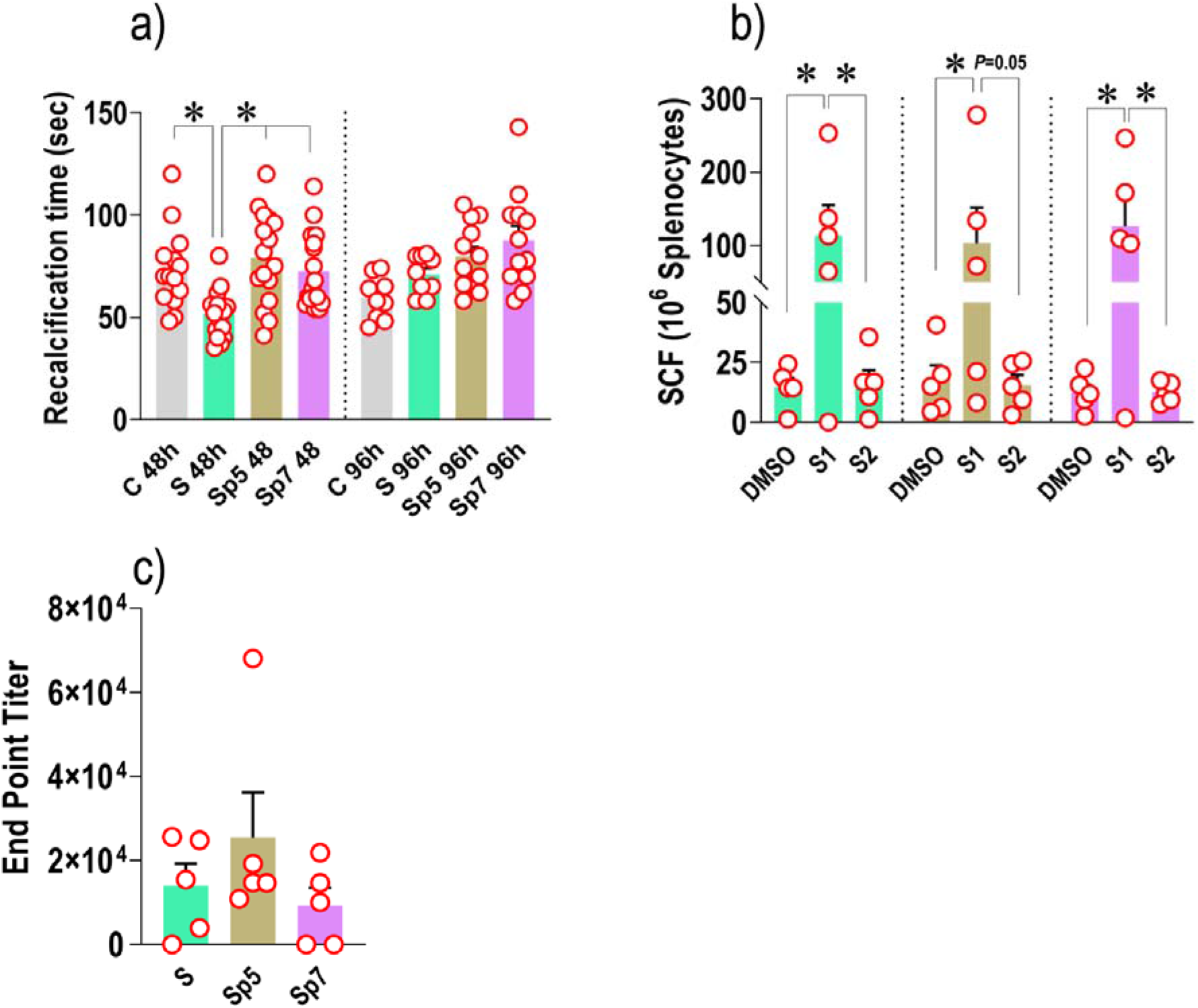
*In vivo* pro-coagulation activity and immunogenicity of wild-type S-protein and Sp5/Sp7 mutants. Panel **a** indicates the pro-coagulation activity (expressed as a recalcification time) of the serum of mice inoculated with vectors carrying the wild-type S-protein protein and Sp5/Sp7 mutants. The coagulation time of wild-type S-protein receiving mice was decreased by ~30% at 48hrs after vector electroporation, indicating a higher propensity of the blood to coagulate. This phenomenon was transient, as shown by the return to basal level of the recalcification time at 96hrs post-inoculation. Panel **b** indicates the cell-mediated immunity in mice inoculated with vectors carrying the wild-type S-protein and Sp5/Sp7 mutants. The graph shows the measure of IFN-γ-producing T cells by ELISpot assay performed on splenocytes collected from mice at 35 days post-vector electroporation. Panel **c** indicates the levels of anti-S-protein IgGs measured in sera of mice inoculated with vectors carrying the wild-type S-protein and Sp5/Sp7 mutants (prime-boost regimen) at 35 days post-vector electroporation. No differences between groups were observed in the two assays suggesting a similar ability of the Sp5 and Sp7 mutants to elicit a cell-mediated and an antibody mediated immune response. * in **a** and **c** indicate *P* < 0.05 as calculated by one-way ANOVA with Tukey multiple comparisons post-*hoc* tests. * in **b** indicate *P* < 0.05 as calculated by repeated measures one-way ANOVA with Tukey multiple comparisons post-*hoc* tests The number of experimental replicates included in the analysis it is indicated by the individual circles overlapped to graphs, each representing an independent animal.

The previous results suggest the relevance of the interaction between S-protein and ERα for the activation of the coagulation cascade *in vivo*. They, however, did not clarify whether the two mutated versions of S-protein elicit a similar cell-mediated immune response and production of antibodies against SARS-CoV-2. This issue is very important for the translation of second-generation RNA/DNA vaccines with a minimized risk of thrombogenicity (*11*). In order to compare the ability of the two mutated proteins to elicit a cell-mediated response and antibodies production, we performed an ELISPOT analysis using splenocytes derived from mice receiving the wild-type and the two mutant proteins, and titrated the circulating anti-S-protein antibodies, both at 35 days after vectors electroporation. Results shown in **Figures 4b, c** demonstrate that both the Sp5 and Sp7 proteins are immunogenic and that they are similarly efficient in inducing cell-mediated immune response against the S1 region of SARS-CoV-2 and the production of antibodies.

## Conclusions

The data contained in the present report provide a preliminary indication that a contributing cause to the hypercoagulatory syndrome present in patients with COVID-19 could derive from an infection-independent activity of the SARS-CoV-2 S-protein mediated by crosstalk with the transcriptional pathway controlled by ER-α (Solis et al., *Sci. Adv*. in press), and resulting into procoagulant activity and TF up-regulation. This hypothesis is supported by the dose-dependent increase in the PCA and *TF* mRNA expression in endothelial cells subjected to stimulation with S-protein and its inhibition by antagonists of the ERα with SERMs and SERDs characteristics (*12*) (**Figure 1**). It is further corroborated by the evidence that the two mutant versions of S-protein (Sp5 and Sp7, **Figure 2**) designed to not to interact with ERα, increased the PCA and *TF* expression/activity in ECs at lower levels compared to the wild-type version of the protein, and did not accelerate coagulation *in vivo* (**Figure 3, 4a**). Given the preliminary nature of these observations we cannot conclude whether the putative S-protein/ERα complex has the ability to directly bind to specific regulatory sequences in *TF* as well as other coagulation factors in ECs or whether it affects other transcriptional circuits involved in the control of the coagulation cascade; we are investigating this currently with a multi-level approach.

In summary, also corroborated by mounting evidences from clinical (*30, 34–36*) and *in vitro* studies (*37*) showing the potential effectiveness of SERMs (e.g. raloxifene) as an anti-viral agent in COVID-19 and other infectious diseases, our results suggest a new non-infective pathologic action of the S-protein at the vascular level, where it has been suggested already to mediate inflammation and vascular dysfunction (*28*). We finally show that Sp5 and Sp7 S-protein versions may be used as a basis to design new versions of DNA/RNA COVID-19 vaccines lacking any residual pro-coagulatory action which, in extremely rare cases (*10*), may cause risk for thrombosis despite the overall very low level of circulating S-protein detected after immunization (*29*).

## Notes

### Competing Interest Statement

FC, MM, CT, DI, FG, ARB and MA serve as employees of the private company Dompe Farmaceutici S.p.A that owns IP rights related to the study findings reported in this paper.

